# Timing and social dynamics of divorce in wild great tits: a phenomenological approach

**DOI:** 10.1101/2024.12.13.628365

**Authors:** A.D. Abraham, B.C. Sheldon, J.A. Firth

## Abstract

Social behaviour is a key part of life for many species. In monogamously breeding species, social associations between breeding partners are particularly important. Selection of a breeding partner often begins well before reproduction, and this process can affect subsequent reproductive success. Thus, the non-breeding season can shape behaviour during the breeding season. However, it is currently unknown how breeding season outcomes can impact associate choice during the non-breeding season, as studying this requires high volumes of cross-context individual social data. This study used three years of wild great tit social data from Wytham Woods, Oxford, UK, to examine social associations between pairs classified with respect to prior and future breeding status (divorcing, faithful, new and juvenile). We found distinct patterns of social association in ‘divorcing’ pairs from early winter, suggesting that divorce is an ongoing behavioural process. Newly forming pairs initially associated similarly to divorcing pairs, but became similar to faithful pairs over time. On a finer spatiotemporal scale, the behaviour of faithful and divorcing birds diverged over the winter. These results provide the first evidence of a behavioural signature of divorce during the non-breeding season in great tits, while suggesting that different behaviours may drive behavioural divorce at different times.

## 4 Introduction

For many animals the social environment is a source of constant change, forcing individuals to make a variety of social decisions (Kurvers et al., 2014). The number and identity of social associations of an individual may have wide-ranging fitness consequences for processes such as predation mortality (Alberts, 2019; Kelley et al. 2011; Montero et al., 2020), disease spread (Behringer et al., 2006; Evans et al., 2020; Stroeymeyt et al., 2018), and foraging activity (Lachlan et al., 1998; Machado et al., 2019). Research into these social processes has often defined sociality on the group level, for example by summarising an individual’s associations across the population. However, the identity of those associations is another important aspect of social behaviour. For example, individuals may be more likely to have social relationships with kin (Podgórski et al., 2014; Wilkinson et al., 2019), and long-term associations with specific individuals may have positive fitness impacts (Emery et al., 2007; Griffiths et al., 2004; Silk et al., 2009).

Mate choice is a particularly clear example of an important individual-level social choice, particularly in socially monogamous species. The potential fitness consequences of mate choice have been widely discussed (Jennions and Petrie, 1997; Kokko et al., 2003), and many studies have shown that remating with a previous partner (pair fidelity) may have consequences for reproductive success (Black, 2001; Gokcekus 2023; Sánchez-Macouzet et al., 2014). However, this key process of selecting a mate is not only relevant during the breeding season. Indeed, social interaction with potential partners in a non-breeding period has been shown to influence mate choice (Cheetham et al., 2008; Senar et al., 2013; Yokoi et al., 2016) and familiarity with a mate in the non-breeding season may increase pair quality and reproductive success (Culina et al., 2020; Griggio and Hoi, 2011; Maldonado-Chaparro et al., 2021; Martin and Shepherdson, 2012). These examples provide evidence that individual social associations are relevant across the breeding and non-breeding contexts, for example in great tits (*Parus major*), who have higher fitness when in a pair that met earlier in the non-breeding season (Culina et al., 2020). Despite this, how key breeding partner bonds persist after the breeding season, or if they dissolve during the non-breeding season, remains largely unknown in many monogamous species.

In animals that mate across multiple seasons, a tradeoff may exist between the benefits of remaining with a familiar partner and the costs of remaining with a less optimal one. Several studies show that increased familiarity with a partner (i.e number of years having bred together) is associated with increased reproductive success (Gokcekus et al., 2023; Schieck and Hannon, 1998). At the same time, the chance of an individual divorcing (separating from its previous partner and repairing with a new partner while its previous partner is still alive) often increases when a pair’s reproduction is less successful (Choudhury, 1995; Culina 2014; Dubois and Cézilly 2002; Linden, 1991). There is limited research into how, in iterated breeding attempts, changes in social behaviour between years maintain these ongoing relationships, particularly in social contexts outside of the breeding season. If the dyadic social behaviour of pairs which go on to be either faithful or divorced can be distinguished in the non-breeding season immediately after a breeding attempt, that suggests that individuals may adjust their social associations in response to information from the breeding season.

The social structure of great tits (*Parus major*) provides an opportunity to study this transition and interaction between breeding and non-breeding sociality. In the winter, great tits feed in loose fission-fusion flocks as they move around the woodland (Aplin et al., 2012). In the spring, they are territorial and socially monogamous, feeding and caring for a brood in a pair (Patrick et al., 2012). Further, there is much individual-level variation in pair bonding behaviour (Firth et al., 2018), and the timing of pairing appears to relate to breeding success (Culina et al., 2020). These findings suggest that there may be a fitness component to the transition between fission-fusion and monogamous sociality in great tits. This makes them a valuable species in which to study questions about the social processes of pair bond formation and dissolution.

In this research we combine two datasets from great tits in Wytham Wood, Oxford, UK. In the non-breeding season, social associations were inferred using visits of radio-frequency identification (RFID) tags to feeders. Paired birds were then observed during the breeding season, matching identities between the winter and spring. The matched identities of individuals between seasons provided a high volume of individual social data across seasons, allowing us to investigate the connection between spring and winter social behaviour. We focus on how pair retention in the spring relates to sociality between that pair in the winter, within their broader social context. This provides new insights into divorce and partner choice in the wild, and a foundation for understanding how individual social associations transition across social structures and seasons.

## 5 Methods

### 5.1 Data collection

The data for this analysis comes from a long-term study of great tits in Wytham Woods, Oxford, UK (Dryad DOI: 10.5061/dryad.vq83bk453; Abraham, Firth, and Sheldon 2025). Each spring, the tits breed almost exclusively in artificial nestboxes placed throughout the woods. These nestboxes are checked regularly (see Perrins, 1965) and breeding individuals identified using British Trust for Ornithology rings and PIT (passive integrated transponder) tags. Within the five years of breeding attempt data used in this analysis (2011-2015), both parents were successfully identified in 61.9% of nests with any eggs laid (1,330 of 2,148), and 85.4% of nests which produced fledglings (1,185 of 1,388)

During the winter season, 65 supplementary feeders were placed throughout the woods at approximately 250 m even grid spacing. For 13 weekends (Saturday and Sunday only) across three winter seasons (3 December 2011 to 26 February 2012; 1 December 2012 to 24 February 2013; 30 November 2013 to 23 February 2014). Supplementary feeders were only run in the weekend (2 days of every 7). They used RFID (radio frequency identification) antennae to identify bird PIT tags, recording each visit of tagged birds to the feeder. Feeders scanned for PIT tags every one-third of a second to obtain this data. Flocking events were identified from the feeder visit records using a Gaussian mixed model through the asnipe package (Farine, 2013), where clusters of visits to the feeder are identified as part of the same event. For a detailed description of this methodology, see Farine et al., 2015.

### 5.2 Data preparation

All data preparation was done in R (Version 4.2.2; R Team, 2022) using the tidyverse (Version 2.0.0; Wickham et al., 2019) and data table (Version 1.14.8; Dowle and Srinivasan, 2023) packages.

Pairs were defined by their relationship to three time periods. Individuals were first identified if they were present at the feeders during the winter (t). Their breeding partner in the previous spring (t-1) and in the following spring (t+1) was used to identify breeding pairs relevant to our analysis. Pairs were only retained if both individuals were present once or more during the focal winter (t). In total, there were six pair statuses which pairs could be assigned, but only four were used in our analysis (Figure 1; see Appendix (A.1) for details of unused pair statuses and the pair categorisation process):

**Figure 1:**
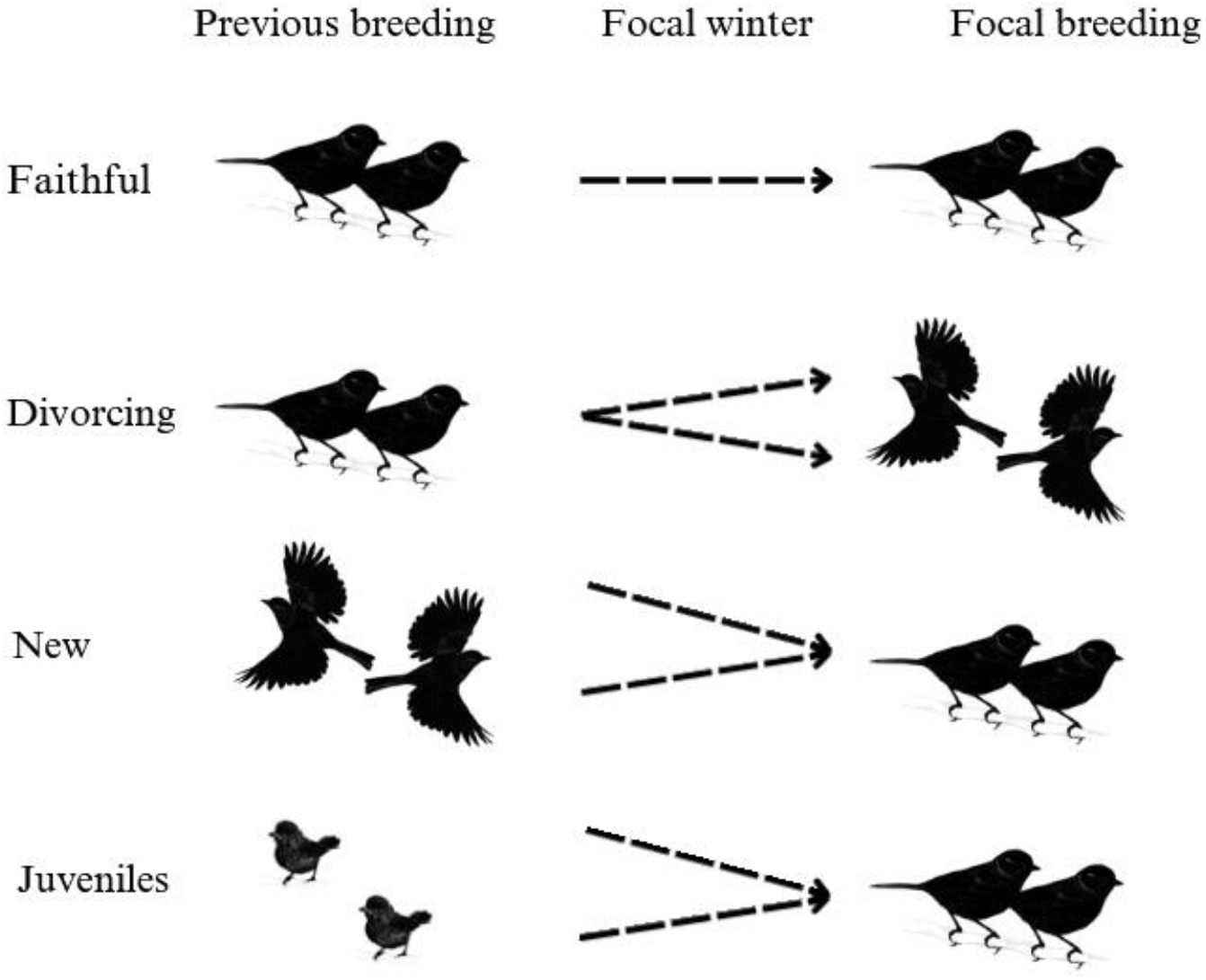
The relationship between pair status in subsequent breedings and pair status as used in the analysis. Focal winter refers to the winter in which behavioural indices for the pair are calculated.

#### Faithful

The pair was observed together in the previous (t-1) and focal (t+1) breeding.

#### Divorcing

The pair was observed together in the previous (t-1) breeding, and observed with other individuals in the focal breeding (t+1) (or either individual was not observed in the focal spring (t+1), but observed alive at a later date, and therefore known to not have died, while their partner was observed with a different individual (t+1)).

These pairs are referred to as ‘divorcing’, despite their divorced status being known at t+1, to ensure that they are distinguished from new pairs formed of two divorced birds. While birds that are not observed as re-paired may still be divorced, only pairs where one or both birds are known to have re-paired (and if only one observed re-paired, the other known still to be alive) are classified as divorced for this analysis, to ensure their divorced status is known. For a discussion of identifying pair fidelity in this system, see Culina et al. 2013. We have used the term divorced for continuity of terminology with literature in this system (Culina, Hinde, and Sheldon 2015) and others (Choudhury 1995; Jeschke and Kokko 2008; Speelman 2024).

#### New

Each member of the pair was observed with other individuals in the previous breeding (t-1), and observed together in the focal breeding (t+1). Alternatively, one pair member was observed with another individual in the previous breeding (t-1), and the other pair member was a juvenile.

There is overlap between ‘Divorcing’ and ‘New’ pairs, with an individual able to be recorded twice in the same year, once as part of a ‘Divorcing’ pair, and once as part of a ‘New’ pair. Across the pairs that were used for analysis (see Table 1), 27 individuals were observed as part of two pairs in the same year.

**Table 1:** The number of pairs in each pair status, across the three seasons, which were suitable for analysis. Pairs in 2012 were observed in the 2012 breeding season, and analysed using data from the 2011/2012 winter season, and so on for the following years.

| Status/Year | 2012 | 2013 | 2014 | Total |
| --- | --- | --- | --- | --- |
| Faithful | 23 | 27 | 27 | 77 |
| Divorcing | 9 | 12 | 5 | 26 |
| New | 30 | 29 | 34 | 93 |
| Juvenile | 44 | 8 | 39 | 91 |
| **Total** | **106** | **76** | **105** | **287** |

#### Juveniles

Both individuals in the pair were born the previous year (t-1), and the focal breeding (t+1) was their first breeding season.

### 5.3 Data analysis

Generalised linear mixed models were run using the glmmTMB package in R (Brooks et al., 2017), with residuals checked used the DHARMa package (Hartig, 2022). Where models failed visual residual assumption checks, they were further tested using non-parametric permutations, to also obtain permuted p-values for the results (p_perm_). Details of model results, residual checks, and further discussion of permutation testing, are included in the Appendix. Each of the three basic indices (see below) were calculated for each pair for every day both individuals were present within the season.

As data was only collected on weekends, and the difference in start dates between the years is negligible, day is coded as ‘experimental day’, where 1 represents the first day of data collection for a given year, to allow seasonal measures to align across years. The experimental day variable represents the number of weekdays and weekends (i.e the second weekend was experimental days 8 and 9), in order to more accurately represent the scale of change over the season. Where a pair wasn’t present on a given day, the value of the behavioural indices for that day would be recorded as NA, and therefore not included in the model.

### Shared flocking events

In order to estimate social associations we examined how individuals co-occurred across flocking events, following the gambit-of-the-group approach (Franks et al., 2010). For each pair, a winter association score (WA) was calculating using the daily proportion of shared flocking events (also known as simple ratio index (SRI)) using the following formula:

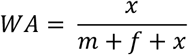

Where *x i*s the number of shared flocking events between a pair, *m* is the number of flocking events observed to include the male but not the female, and *f* is the number of flocking events observed to include the female but not the male.

Winter association score was calculated and modelled at the pair level (i.e one WA for each pair). A k/n binomial generalized linear model was used. In order to fit the score to the model structure, the response variable was coded as successes=*x* and failures=*m+f+x*, making the response WA. Pair status was the fixed effect predictor. Year and pair ID were included as random effects. Pair ID was a combination of the IDs of the two individuals in a pair, meaning pairs which appeared multiple times (i.e faithful pairs) had the same pair ID across years. A 1:n overdispersion random effect and a zero-inflation term were included, as determined by AICc model selection (Table A2).

To model WA over the course of the season, the additional effect of experimental day was calculated by including a quadratic interaction term between pair status and experimental day, as a linear interaction model did not converge. All other fixed and random effects were the same as in the aggregate model.

### Preferred partner

In order to investigate whether the behavioural differences we observed reflected a difference in general social behaviour or a specific difference in behaviour with an individual’s partner, we analysed whether individuals’ breeding season partner remained their ‘preferred social partner’ in the winter season. A preferred social partner was defined as the opposite-sex individual with the highest winter association score with the focal individual. In this dataset, of birds with closest associates of known sex, 70% had a closest associate of the opposite sex (73% of females and 67% of males). In cases of the closest associate being of unknown sex, the closest associate known to be of the opposite sex was identified as the preferred social partner.

Unlike winter association score, partner preference was not symmetrical. That is, each individual in a pair may have a different result for whether their preferred social partner was their breeding partner. Therefore, preferred partner was modelled on the individual level not the pair level. Pair ID was still included as a random effect to account for non-independence within the pair.

Modelling partner preference aggregated across the winter was not possible with a binary logistic regression (1 = preferred partner same as breeding partner, 0 = preferred partner different to breeding partner) due to perfect separation of the divorced category (no divorced birds had their breeding partner as their preferred social partner). Due to the clear division of the data, no modelling was undertaken, and the more complex model of preferred partner over time was used to infer overall differences between the pair statuses.

In the model of preferred partner over time, the response was a binary variable, where 1 = preferred partner the same as the breeding partner, and 0 = preferred partner different to breeding partner. This was modelled using a logistic regression. Pair status in interaction with a quadratic experimental day term and sex were used as fixed effects. Year and pair ID were included as random effects.This model was compared to a linear interaction model using AICc, and the quadratic model performed better so was retained (Table A12).

### Visit adjacency

We also examined pairs’ behaviour within flocking events. Daily visit adjacency was calculated only from records within flocking events that pairs shared. For each record, it was marked as adjacent if an individual visited the feeder within 3 seconds after their partner. This measure was investigated in order to investigate co-occurrence on a smaller spatial scale than shared ratio index, which measures only co-occurrence in flocking events that may be made up of many individuals over varying lengths of time. The assumption underlying our use of visit adjacency was that visiting the feeder within quick succession would represent a prioritization of social contact with a given individual.

The amount of adjacent visits by a pair was estimated as a simple proportion, referred to from here as visit adjacency index (VAI):

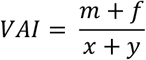

where *m* is the number of male visit records adjacent to a female visit, *f* is the number of female visit records adjacent to a male visit, *x* and *y* are the total number of male and female visits respectively, and *VAI* is the visit adjacency index.

Aggregate VAI across the winter was modelled using a k/n binomial generalised linear model. The response was coded as successes (k) = m+f and failures (n) = x+y, making the response equivalent to VAI. Pair status was included as a fixed effect, and year and pair ID as intercept-level random effects. Average flock size was also included as a fixed effect, to account for follower proportion presumably being lower in larger flocks. A 1:n overdispersion random effect was included, as selected by AICc (Table A4).

The model of visit adjacency over time was the same as the base model, but a quadratic interaction term was included between pair status and experimental day. The model was compared to a linearly modelled interaction using AICc, but the quadratic model performed better so was retained.

### Non-pair associates

As an additional analysis we compared the winter association and visit adjacency scores of birds with their partner to the same scores with their other network associates. This was in order to understand whether any results observed might be due to a change in behaviour with all conspecifics, as opposed to just with the individual’s partner. Details of model building and results for this analysis are included in A.3.

## 6 Results

Of 1,169 breeding pairs observed across three seasons, the total number of pairs which fulfilled the criteria for this analysis was 287 (24.6%). 635 pairs (54.3%) were ineligible because one or both individuals were not observed at the feeders in the focal winter, meaning no data was available for analysis, and 247 (21.1%) pairs had unclear pair status and were therefore removed (Table A1). The number of analysed pairs of each status in each breeding season is presented in Table 1. A relatively low number of eligible pairs is expected because of the high mortality and potential for immigration in this wild system. A full breakdown of the workflow for which pairs were and were not included is provided in A.1.

### 6.1 Shared flocking events

In a model of winter association score aggregated across time, there was a significant difference between pair statuses (Table A3). With faithful as a reference value, new (coef=-0.54*±*0.15, p*<*0.001, p_perm_=0), juvenile (coef =-0.88*±*0.15, p*<*0.001, p_perm_=0.01), and divorcing (coef=-1.31*±*0.23, p*<*0.001, p_perm_=0) pairs all had significantly lower winter association. Faithful birds had significantly higher winter association score with their partner than with non-pair associates (coef=-1.08*±*0.04, p*<*0.001 (Table A13)), while divorcing birds had similar winter association with their partner when compared to their non-pair associates (Figure A10).

Across experimental days, faithful pairs showed a significant increase in winter association score (coef=0.24*±*0.02, p*<*0.001), with no significant quadratic effect (Figure 2). New pairs had a significant linear increase in winter association over time (0.14*±*0.03, p*<*0.001), with an n-shaped quadratic curve (−0.09*±*0.03, p=0.006). Juvenile pairs also had a significant linear increase (0.29*±*0.04, p*<*0.001), but in contrast, a u-shaped curve (0.09*±*0.04, p=0.01). Divorcing pairs had a significant linear decrease in winter association score over time (−0.26*±*0.05, p*<*0.01), with no significant quadratic effect. A full model coefficient table is provided in the Appendix (Table A7).

**Figure 2:**
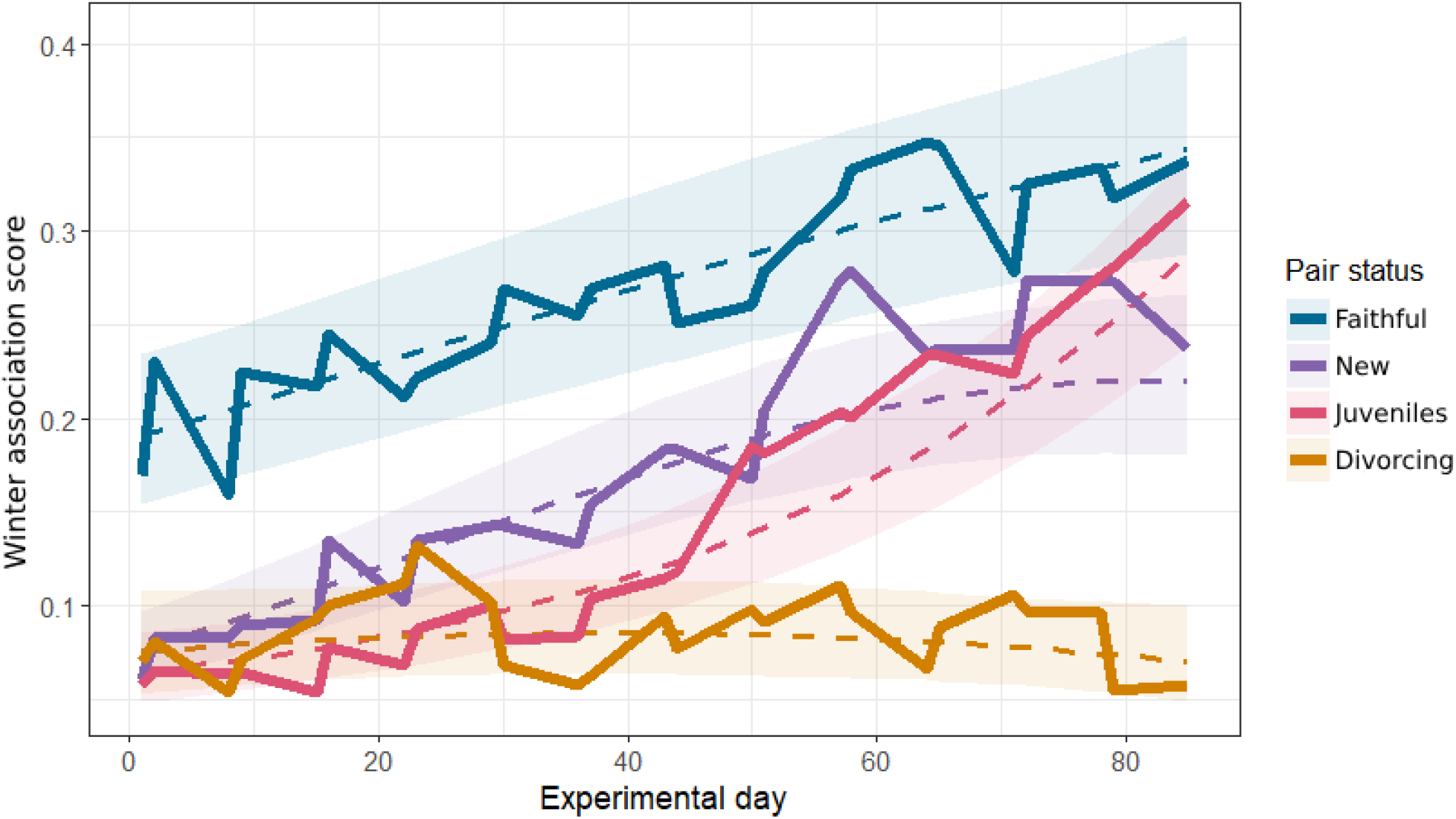
The predicted and observed association between experimental day and winter association score. Dashed lines represent the predicted value from a binomial GLMM with a quadratic interaction term, and ribbons a 95% confidence interval around that prediction. Predictions are made for each pair status at each relevant value of experimental day, averaged across random effect groups and conditioned on the zero-inflation term (Lüdecke, 2018). Solid lines are the average values from the raw data.

### 6.2 Preferred partner

As discussed in the methods, we could not model the effect of pair status on the likelihood of a breeding partner being a preferred partner, due to perfect separation of the divorcing category. However, the existence of perfect separation suggests the difference may be significant, and the overall effect can be extracted from the model of preferred partner over time.

Aggregated across the season, new (coef=-0.76*±*0.17, p*<*0.001), juvenile (−1.09*±*0.18, p*<*0.001), and divorcing (−2.84*±*0.41, p*<*0.001) individuals were significantly less likely than faithful individuals to have the highest association with their breeding partner (Figure 3). Faithful individuals had a significantly increased likelihood over experimental day (0.38*±*0.05, p*<*0.001), with their rate of increase slightly decreasing towards the end of the winter (−0.13*±*0.06, p=0.02). Individuals in new pairs had a significant linear increase over time (0.27*±*0.09, p=0.002) and a similar decrease in rate, causing an n-shaped curve (−0.23*±*0.09, p=0.01). Individuals in juvenile pairs significantly increased the chance that their partner was their highest association over time (0.71*±*0.11, p*<*0.001), with no significant quadratic effect. Divorcing individuals were significantly less likely to have the highest association with their partner over time (−0.67*±*0.23, p=0.003), but with no significant quadratic effect. There was no effect of sex. A full model coefficient table is provided in the Appendix (Table A12).

**Figure 3:**
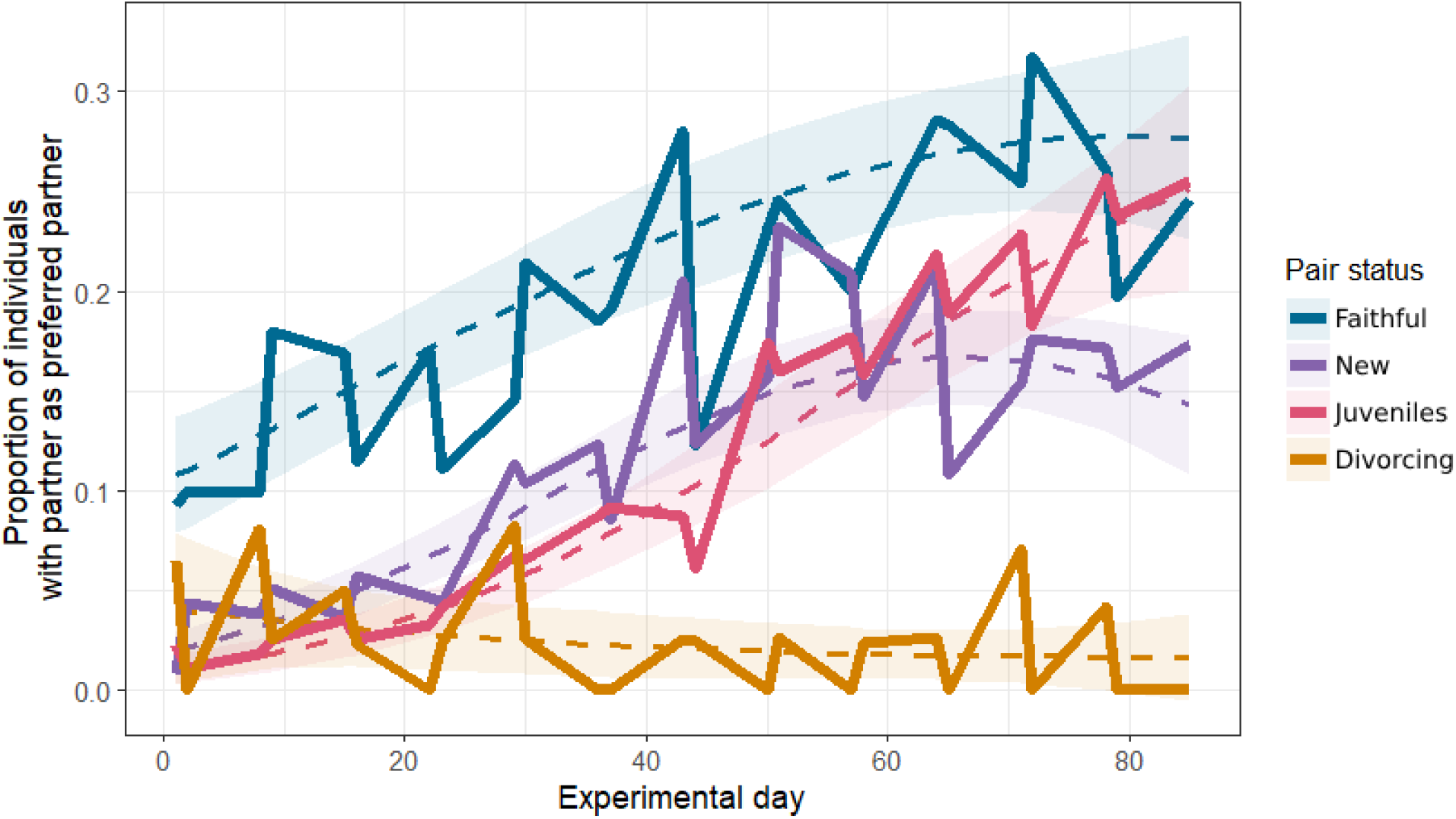
The predicted and observed association between experimental day and whether an individual had the highest winter association score with their breeding partner. Dashed lines represent the predicted value from a logistic GLMM with a quadratic interaction term, and ribbons a 95% confidence interval around that prediction. Predictions are made for each pair status at each relevant value of experimental day, averaged across random effect groups and conditioned on the zero-inflation term (Lüdecke, 2018). Solid lines are the average values from the raw data.

### 6.3 Visit adjacency

Examining behaviour within flocking events, we found no significant difference between overall visit adjacency index for faithful, new (coef=0.05±0.06, p=0.42), and juvenile (0.08±0.07, p=0.28) pairs. However, divorcing pairs had significantly lower visit adjacency index (−0.54±0.12, p<0.001) (Table A5). Faithful birds had significantly visit adjacency index with their partner than with non-pair associates (coef=-0.42±0.04, p=<0.001 (Table A14)), while divorcing birds had similar visit adjacency index with their partner when compared to their non-pair associates (Figure A13)

In a quadratic interaction model between pair status and time, there was a slight but significant linear increase in visit adjacency index over the winter for faithful (coef=0.05±0.02, p=0.03) and juvenile (0.10±0.04, p=0.01) pairs (Figure 4). There was a significant linear decrease in visit adjacency index for divorcing pairs (−0.23±0.08, p=0.006). New pairs did not have a significant linear change over time (0.06±0.04, p=0.1). Despite the quadratic interaction model outperforming a linear model, none of the quadratic effects in the model were significant. The conclusions drawn from the linear interaction model coefficients were the same as the quadratic model. Full model coefficient tables (quadratic and linear model results) are provided in the Appendix (Table A9, Table A10).

**Figure 4:**
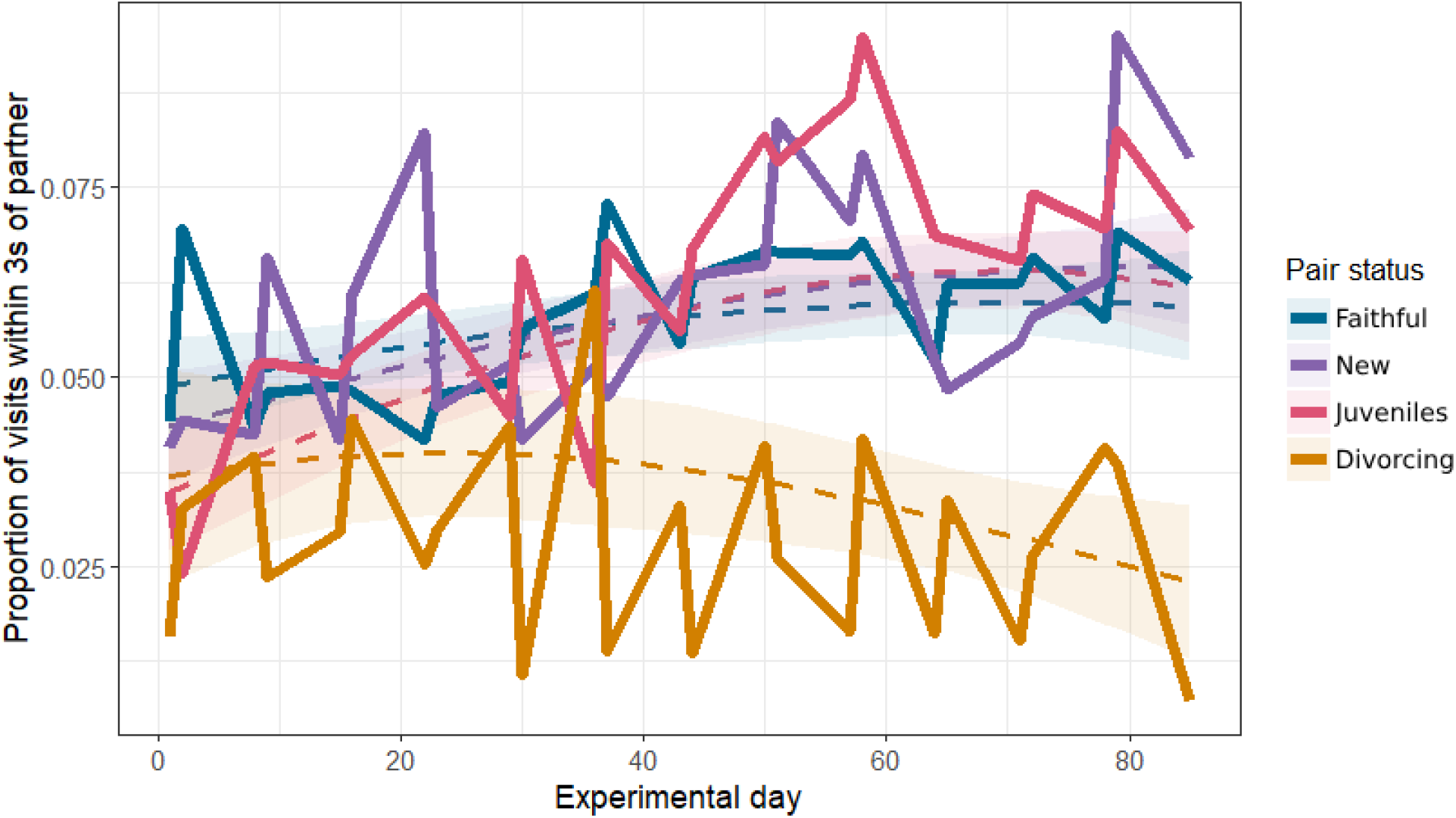
The predicted and observed association between experimental day and visit adjacency index. Dashed lines represent the predicted value from a binomial GLMM with a quadratic interaction term, and ribbons a 95% confidence interval around that prediction. Predictions are made for each pair status at each relevant value of experimental day, averaged across random effect groups and conditioned on the zero-inflation term (Lüdecke, 2018). Solid lines are the average values from the raw data.

## 7 Discussion

We show how socially monogamous breeding relationships carry over into behaviour in a group context, and find that social behaviour during the non-breeding winter season is strongly associated with pair status in the neighbouring springs in a wild population of great tits. Specifically, we report clear evidence that social associations among pairs undergoing the process of divorce exhibit very different patterns compared to pairs that remain faithful or new pairs that are forming., We showed that across the winter as a whole, faithful pairs of wild great tits shared the most flocking events with their breeding partners, and divorcing pairs the least. These differences are present in both absolute winter association score between a pair, and the chance of an individual having the highest winter association score with their breeding partner, suggesting an active adjustment of time spent with a partner, as opposed to a change in behaviour with all conspecifics. Within those shared events, pairs which were together in the subsequent spring (faithful, new, and juvenile) had considerably more visits to the feeder immediately after their partner, when compared to pairs which were divorcing.

The distinction between faithful and divorcing pairs from the very beginning of the winter period provides new insights suggesting that information from the breeding season almost immediately influences individual social associations during the winter non-breeding season.. However, the different pattern observed in visit adjacency means that there may be causal differences underlying the results at different spatiotemporal levels of behaviour. Shared flocking events are inherently spatial, as it is impossible for birds to share a flocking event if they are not in the same location. It is uncertain to what extent shared flocking events reflect social decision making as opposed to simply non-deliberate spatial co-occurrence (i.e high winter association score between two birds could be due to them remaining in the same area of the woods, as opposed to choosing to associate together). Previous work in this system has suggested that large-scale social structure of the great tits is explained to some degree by spatial structuring, but that these spatial effects cannot explain all patterns in the tit social networks (Aplin et al. 2015; Farine and Sheldon, 2019). Pair closeness within flocking events may reflect a more direct social decision at a fine spatiotemporal scale, therefore representing social behaviour over and above spatiotemporal location. It may be that at the beginning of the season the difference between faithful pairs and other pairs is spatially determined, while social decision-making becomes more important throughout the season. This may represent a general insight into the processes that determine pair bonding and fidelity. To separate social behaviour from spatial feeding behaviour, further experiments which separate individual feeding locations during the non-breeding season are now needed (see Firth and Sheldon, 2015 for an example of a feeder segregation experiment).

Previous work in this system has shown associations in the winter carrying into the breeding season, in both non-breeding social associates (Firth and Sheldon 2016) and breeding pairs (Culina et al 2020). The results provided here ‘complete the circle’, helping to contribute to a full picture of how social associations carry between contexts. They further suggest an important role of the individual identity of associates. The great tits in our study showed a difference in their social behaviour with a previous partner related to whether they would go on to breed together again. A potential explanation for this difference could be that birds which are more likely to divorce are birds which already have some distinct pattern of social behaviour, such as having a less strong pair bond with their partner even before breeding (Culina, Hinde, and Sheldon, 2015). Alternatively, it could be that divorce leads to a change in overall behaviour (this has been shown in humans (Milardo, 1987)), not just behaviour with a previous partner, such as an increased number of weaker social bonds as an individual searches for a new partner. There may also be a selective disappearance effect in this data. Previous research has found that pairs which separate have lower survival (Culina 2013). This will, therefore, selectively remove a group of divorcing birds from the pairs used in analysis, which could obscure true patterns in behaviour and the non-behavioural consequences of divorce. Further research into the underlying behavioural processes may help to disentangle these potential explanations.

The results of this analysis have potentially wider implications for questions in the analysis of dynamic network systems. In particular, we have shown that temporal patterns in dyadic features in this network (e.g winter association score) could be used to predict states at another time (e.g divorce). Regardless of the causality of these patterns, the predictive element could be used in analogous ecological and epidemiological networks. There is room for further study of this dynamic, particularly in the context of how relationships in this network evolve along the cyclical seasonal gradient.

In sum, we provide new insights into the process of divorce and new partnership formation in a wild population, and demonstrate how social relationships during the breeding season shape non-breeding winter and social decisions in great tits. To our knowledge, this is the first work identifying an associate-level, non-breeding signal of divorce using distinct behavioural patterns. Future examinations focusing on investigating the causal relationships behind the observed associations in this and broader systems would now be of much interest.

## 1 Acknowledgments

We would like to thank the many people who have helped to collect data used in this research: those who collected long-term breeding data from the Wytham tit study, and members of the EGI Social Network group who collected social network data and developed methodology for studying the social networks of great tits. Thank you to two anonymous reviewers for their helpful comments on the manuscript. The long-term population study used here for breeding data has been supported by grants including BBSRC (BB/L006081/1) and NERC (NE/K006274/1, NE/S010335/1); the social network data were funded by NERC grant (AdG250164). AD Abraham is supported by the Newton-Abraham studentship (no relation!)

