## Appendix for "Timing and social dynamics of divorce in wild great tits: a phenomenological approach"

**A.1	Pair statuses**

Pair statuses for a given breeding season were calculated based on the following variables:

1. Whether the pair was observed during the focal breeding

2. Whether the female was observed in the focal breeding

3. Whether the male was observed in the focal breeding 4. Whether the pair was observed in the previous breeding

5. Whether the female was observed in the previous breeding

6. Whether the male was observed in the previous breeding

7. Whether the female is a juvenile (born in the previous breeding season)

8. Whether the male is a juvenile

9. Whether the male was observed alive in any later season

10. Whether the female was observed alive in any later season

The combination of these variables determined the status of the pair. Certain combinations were ‘impossible’ (e.g pair observed in a year but the female not observed) and were used to identify and correct errors in data entry. The full list of pair statuses, based on the above variables, is as follows:

*Faithful*
A pair was ‘faithful’ if they were observed in the focal breeding (1=TRUE), and observed in the previous breeding (4=TRUE).

*Divorcing*
A pair was ‘divorcing’ if the pair was not observed in the focal breeding (1=FALSE), but both individuals were (2&3 = TRUE), and the pair had been observed in the previous breeding (4=TRUE).

Alternatively, if one individual was not observed in the focal breeding (2 XOR 3=FALSE) but they were observed alive at a later date (9 XOR 10=TRUE), they would still be recorded as divorcing.

*New*
A pair was ‘new’ if they were observed in the focal breeding (1=TRUE), not observed in the previous breeding (4=FALSE), but both individuals were observed in the previous breeding (5&6=TRUE). Pairs were also classified as new if one but not both individuals were a juvenile (7 XOR 8=TRUE).

*Juveniles*

A pair were ‘juveniles’ if they were observed in the focal year (1=TRUE) and both individuals were juveniles (7&8=TRUE).

*Split*
A pair was ‘split’ where they were no longer breeding together, but the fate of one or both birds was unknown so it was unclear whether the split was due to divorce or death. This was the case where they were not observed in the focal breeding (1=FALSE) but were observed in the previous breeding (4=TRUE) and one or both individuals were not observed in the focal breeding (2/3=FALSE) and not observed alive at a later date (9/10=FALSE). This category was not included in analysis.

*New (unknown)*

A pair was ‘new unknown’ where they were breeding together, but their status in the previous breeding was unknown, so it was unclear whether they were a faithful or new pair. This was the case where they were observed in the focal breeding (1=TRUE) but were not observed in the previous breeding (4=FALSE) and one or both individuals were not observed in the previous breeding (5/6=FALSE). This category was not included in analysis.

*Table A1: The number of breeding pairs of each status across each of the three study years. As these totals include birds which did not have movement data from the previous winter, and therefore could not be included in the analysis, totals for the faithful, divorcing, new, and juvenile statuses may be higher than in Table 1. Pairs in 2012 were observed in the 2012 breeding season, and analysed using data from the 2011/2012 winter season, and so on for the following years.*

| Status/Year | 2012 | 2013 | 2014 | **Total** |
| --- | --- | --- | --- | --- |
| Faithful | 27 | 29 | 30 | **86** |
| Divorcing | 12 | 16 | 5 | **33** |
| New | 47 | 50 | 52 | **149** |
| Juveniles | 78 | 17 | 84 | **179** |
| Split | 171 | 218 | 118 | **507** |
| New (unknown) | 111 | 58 | 46 | **215** |
| **Total** | **446** | **388** | **335** | **1169** |

Figure A1 provides an example of how individuals may be assigned to different pair statuses. Due to the individually-focused approach, numbers in the diagram may differ from those in other tables, but it should demonstrate how the number of pairs usable in analysis may reduce dramatically.

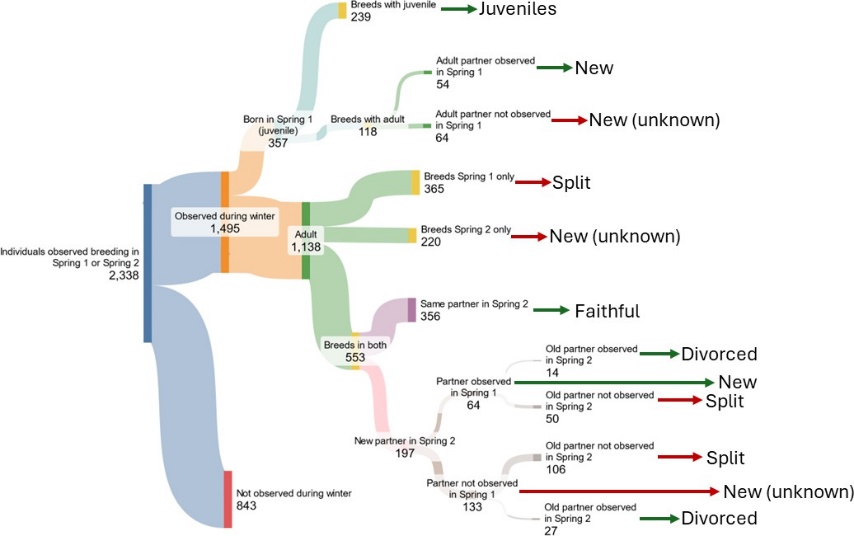

*Figure A1: Number of individuals in the data at each stage of pair classification. Pair statuses denoted with a green arrow are those included in the final analysis, while ones with red arrows were not. Note that as an individual’s partner has to be observed in the winter to be included in the final analysis, numbers may differ from those included in other tables.*

**A.2	Model checking and detailed results**

**A.2.1	Suitability of permutation testing**

In cases where error distributions appear not to follow assumptions of normality, permutation testing is a possible non-parametric solution. It tests whether the observed results differ from the distribution of the values estimated when the data is ‘shuffled’ - if they do, it suggests that any significant relationship found from the original model did not arise by chance. The probability of obtaining the observed value from the permuted distribution provides a new p-value.

While this method is very well accepted for models with a single predictor, it is debated whether it remains suitable for models with multiple predictors, random effects, or repeated samples (Kennedy and Cade, 1996). This is because the shuffling may break relationships within the data, making the randomised dataset not a relevant comparison to the original. There are a number of ways to perform a permutation test, as any combination of parameters can be permuted. Theoretically, permutation of the residuals constrained by model groups would provide the most reliable results (M. Anderson and Braak, 2003; Manly, 2007). However, limitations on permutation from using binomial models with a limited number of constrained permutations (possible combinations within groups) mean that permuting the raw data was the most suitable for my analysis.

Following the procedure from Hope, 1968 and M. J. Anderson and Legendre, 1999, we assessed the results of the permutation from the Z-score of the coefficients, as opposed to the raw coefficient values. An exact permutation test was not possible, as that would require all possible permutations, and therefore 534! at minimum for my models. Instead, we conducted randomisation tests, where a large random sample of possible permutations were used. Where residual assumptions were not met for a model, I used the following procedure, for a k/n binomial model.

1. Randomise k/n pairs across the dataset.
2. Run original model structure using randomised dataset.
3. Extract model coefficient Z-values.
4. Repeat 1-3 5000 times.
5. Compare the absolute distributions of the coefficient Z-values from the permuted models to the coefficient Z-values from the original model, in order to obtain a two-tailed p-value.

In interpreting the results of the permutation tests it is important to note that small changes in how the permutations are performed may change the power of the analysis, and that in some cases permutation tests can still be affected by nonhomogeneity of variance. Therefore, where possible, we still attempted to adjust the initial model in order to fulfil residual distribution assumptions, or make breaches of the residual assumptions as minor as possible.

**A.2.2	Shared flocking events**

*Model selection*

*Table A2: AICc model selection results for a model of simple ratio index by pair status. The model used in analysis is italicised.*

| **Model**  **number** | **Fixed**  **effects** | **Random**  **effects** | **Zero-**  **inflation** | **AICc** | ∆ **AICc** |
| --- | --- | --- | --- | --- | --- |
| **1** | Pair  status | Year +  Pair ID | - | 4397.49 | 1226.81 |
| **2** | Pair  status | Year +  Pair ID + 1:n | - | 3233.64 | 62.96 |
| **3** | Pair  status | Year +  Pair ID | *∼*1 | 3948.10 | 777.42 |
| ***4*** | *Pair*  *status* | *Year +*  *Pair ID*  *+ 1:n* | *∼1* | *3170.68* | *0* |

*Model residuals*

The residuals from the selected model showed significant deviations from assumptions (Figure A2). To investigate these deviations as a potential explanation for observed effects, a permutation test was conducted.

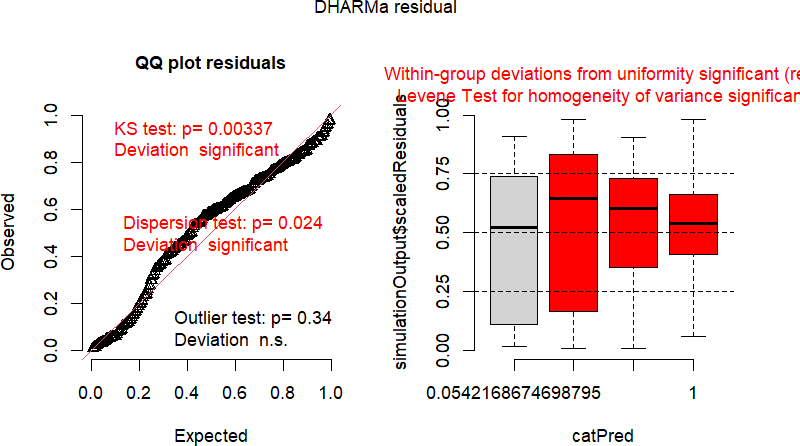

*Figure A2: The model residuals (estimated with DHaRMa) from a binomial GLMM of the effect of pair status on simple ratio index.*

*Permutation test*

A permutation test was conducted, with 5000 randomisations of the response. 5.98% of the models did not converge, meaning a Z-score value was unable to be extracted. The two-sided permutation p-values supported the results from the original model (Figure A3).

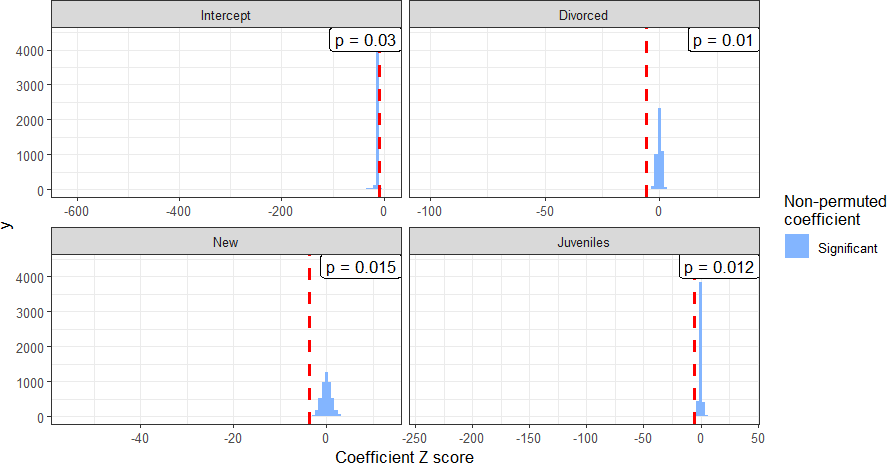

*Figure A3: The result of a permutation test of a model of the effect of pair status on simple ratio index. Non-converged models with NA Z-score values are not included.*

*Coefficient table*

*Table A3: Fixed effect coefficient estimates from a binomial GLMM of the effect of pair status on simple ratio index. Significant coefficients are indicated in italics.*

| **Coefficient** | **Estimate** | **Standard**  **error** | **p-value** | **Permuted**  **p-value** |
| --- | --- | --- | --- | --- |
| *Intercept* | *-1.10* | *0.11* | *<0.001* | *0.03* |
| *Pair status*  *New* | *-0.54* | *0.15* | *<0.001* | *0.015* |
| *Pair status*  *Juveniles* | *-0.88* | *0.15* | *<0.001* | *0.012* |
| *Pair status*  *Divorced* | *-1.31* | *0.23* | *<0.001* | *0.01* |

**A.2.3	Visit adjacency**

*Model selection*

*Table A4: AICc model selection results for a model of visit adjacency index by pair status. The model used in analysis is italicised.*

| **Model**  **number** | **Fixed**  **effects** | **Random**  **effects** | **Zero-**  **inflation** | **AICc** | ∆ **AICc** |
| --- | --- | --- | --- | --- | --- |
| **1** | Pair  status + Average flocksize | Year +  Pair ID | - | 2135.68 | 44.36 |
| ***2*** | *Pair*  *status + Average flocksize* | *Year +*  *Pair ID*  *+ 1:n* | *-* | *2091.31* | *0* |
| **3** | Pair  status + Average flocksize | Year +  Pair ID | *∼*1 | 2137.80 | 46.49 |
| **4** | Pair  status + Average flocksize | Year +  Pair ID + 1:n | *∼*1 | 2093.45 | 46.49 |

*Model residuals*

The model residuals showed no significant deviation from assumptions (Figure A4).

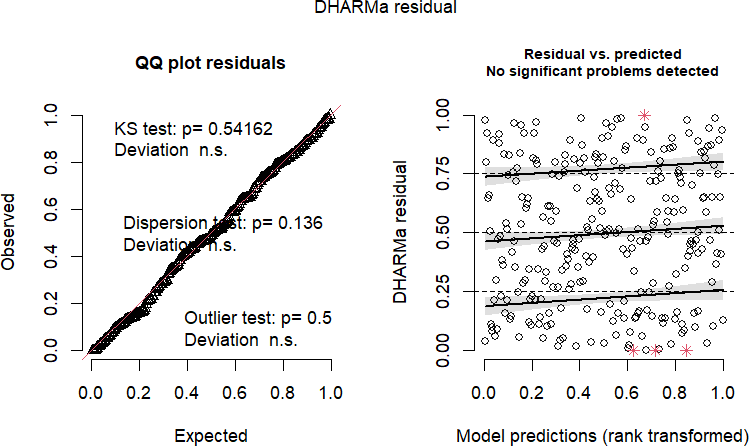

*Figure A4: The model residuals (estimated with DHaRMa) from a binomial GLMM of the effect of pair status on visit adjacency index.*

*Coefficient table*

*Table A5: Fixed effect coefficient estimates from a binomial GLMM of the effect of pair status on visit adjacency index. Significant coefficients are indicated in italics.*

| **Coefficient** | **Estimate** | **Standard**  **error** | **p-value** |
| --- | --- | --- | --- |
| *Intercept* | *-2.45* | *0.09* | *<0.001* |
| Pair status New | 0.05 | 0.06 | 0.416 |
| Pair status  Juveniles | 0.08 | 0.07 | 0.280 |
| *Pair status*  *Divorced* | *-0.54* | *0.12* | *<0.001* |
| *Average*  *flocksize* | *-0.03* | *0.01* | *<0.001* |

**A.2.4	Shared flocking events by time**

*Model selection*

*Table A6: AICc model selection results for a model of winter association score by pair status and experimental day. The model used in analysis is italicised.*

| **Model**  **number** | **Fixed effects** | **Random**  **effects** | **Zero-**  **inflation** | **AICc** | ∆ **AICc** |
| --- | --- | --- | --- | --- | --- |
| **1** | Pair status + Day  + Day^2^ + Pair status*Day + Pair status*Day^2^ | Year +  Pair ID | - | 38036.86 | 9698.97 |
| **2** | Pair status + Day  + Day^2^ + Pair status*Day + Pair status*Day^2^ | Year +  Pair ID + 1:n | - | 29227.10 | 889.21 |
| **3** | Pair status + Day  + Day^2^ + Pair status*Day + Pair status*Day^2^ | Year +  Pair ID | *∼*1 | 30651.73 | 2313.84 |
| ***4*** | *Pair status + Day*  *+ Day*^2^ *+ Pair status*Day + Pair status*Day*^2^ | *Year +*  *Pair ID + 1:n* | *∼1* | *28337.89* | *0* |
| **5** | Pair status + Day  + Pair status*Day | Year +  Pair ID + 1:n | *∼*1 | 28359.61 | 21.73 |

*Model residuals*

The residuals for the selected model show a number of deviations from assumptions (Figure A5). To test these deviations as a potential explanation for the model results, a permutation test was conducted.

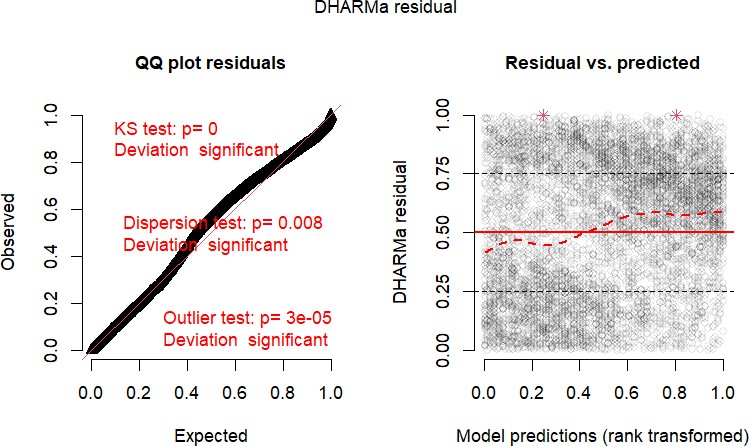

*Figure A5: The model residuals (estimated with DHaRMa) from a binomial GLMM of the effect of pair status and day on winter association score.*

*Permutation test*

A permutation test was conducted, with 5000 randomisations of the response. 3.25% of models did not converge so a Z score was unable to be extracted. The two-sided permutation p-values supported the results from the original model, except for the significance of the Intercept result, with the permutation test suggesting that there may not be statistical evidence that the Intercept value differs from zero (Figure A6). This difference largely doesn’t impact the interpretation of the results.

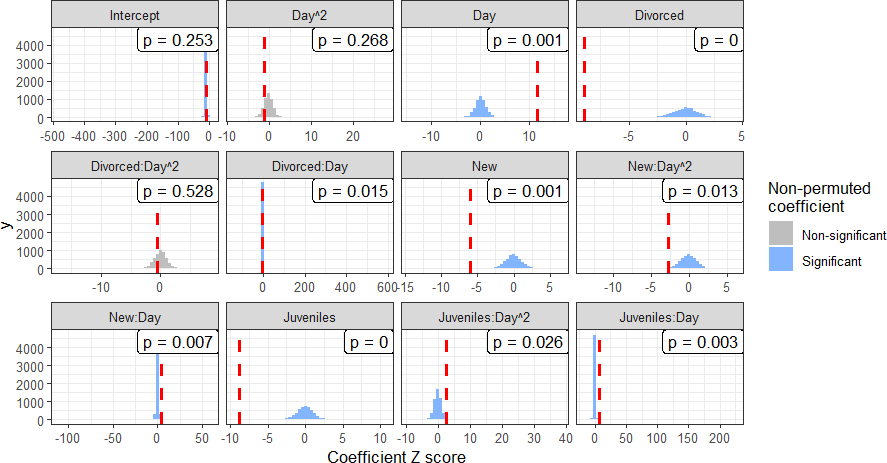

*Figure A6: The result of a permutation test of a model of the effect of pair status and day on winter association score. Non-converged models with NA Z-score values are not included.*

*Coefficient table*

*Table A7: Fixed effect coefficient estimates from a binomial GLMM of the effect of pair status and day on winter association score. Significant coefficients are indicated in italics.*

| **Coefficient** | **Estimate** | **Standard**  **error** | **p-value** | **Permuted**  **p-value** |
| --- | --- | --- | --- | --- |
| *Intercept* | *-0.98* | *0.12* | *<0.001* | *0.253* |
| *Pair status*  *New* | *-0.58* | *0.10* | *<0.001* | *0.001* |
| *Pair status*  *Juveniles* | *-1.01* | *0.11* | *<0.001* | *0* |
| *Pair status*  *Divorced* | *-1.40* | *0.16* | *<0.001* | *0* |
| *Day* | *0.24* | *0.02* | *<0.001* | *0.001* |
| Day^2^ | -0.03 | 0.02 | 0.25 | 0.268 |
| *Pair status*  *New*Day* | *0.14* | *0.03* | *<* 0*.*001 | *0.007* |
| *Pair status*  *Juveniles*Day* | *0.29* | *0.04* | *<0.001* | *0.003* |
| *Pair status*  *Divorced*Day* | *-0.26* | *0.05* | *<0.001* | *0.015* |
| *Pair status*  *New*Day*^2^ | *-0.09* | *0.03* | *0.005* | *0.0013* |
| *Pair status*  *Juveniles*Day*^2^ | *0.09* | *0.04* | *0.014* | *0.026* |
| Pair status  Divorced*Day^2^ | -0.03 | 0.06 | 0.547 | 0.528 |

**A.2.5	Visit adjacency by time**

*Model selection*

*Table A8: AICc model selection results for a model of visit adjacency index by pair status and experimental day. The model used in analysis is italicised. Models which did not converge were not assessed using AICc, so have an AICc value of NA.*

| **Model**  **number** | **Fixed effects** | **Random**  **effects** | **Zero-**  **inflation** | **AICc** | ∆ **AICc** |
| --- | --- | --- | --- | --- | --- |
| **1** | Pair status + Day | Year + | - | 15739.36 | 804.37 |
|  | + Day^2^ + Pair | Pair ID |  |  |  |
|  | status*Day + Pair |  |  |  |  |
|  | status*Day^2^ + |  |  |  |  |
|  | Average flocksize |  |  |  |  |
| ***2*** | *Pair status + Day* | *Year +* | *-* | *14934.99* | *0* |
|  | *+ Day*^2^ *+ Pair* | *Pair ID +* |  |  |  |
|  | *status*Day + Pair* | *1:n* |  |  |  |
|  | *status*Day*^2^ *+* |  |  |  |  |
|  | *Average flocksize* |  |  |  |  |
| **3** | Pair status + Day | Year + | *∼*1 | 15589.54 | 654.55 |
|  | + Day^2^ + Pair | Pair ID |  |  |  |
|  | status*Day + Pair |  |  |  |  |
|  | status*Day^2^ + |  |  |  |  |
|  | Average flocksize |  |  |  |  |
| **4** | Pair status + Day | Year + | *∼*1 | NA | NA |
|  | + Day^2^ + Pair | Pair ID + |  |  |  |
|  | status*Day + Pair | 1:n |  |  |  |
|  | status*Day^2^ + |  |  |  |  |
|  | Average flocksize |  |  |  |  |
| **5** | Pair status + Day | Year + | - | 14938.95 | 3.95 |
|  | + Pair status*Day | Pair ID + |  |  |  |
|  |  | 1:n + |  |  |  |
|  |  | Average |  |  |  |
|  |  | flocksize |  |  |  |

*Model residuals*

Model residuals for the selected model are presented below (Figure A7). There is evidence of over or under dispersion, and patterns in the residuals. While these deviations are relatively minor, a permutation test was conducted to check these residual patterns as a potential explanation for the observed results.

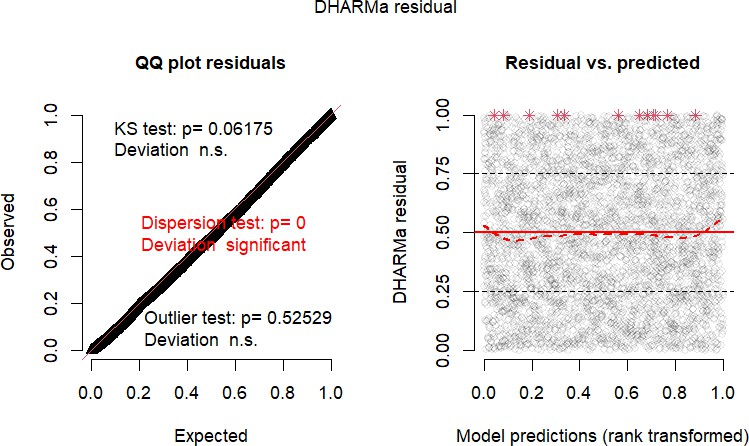

*Figure A7: The model residuals (estimated with DHaRMa) from a binomial GLMM of the effect of pair status and day on visit adjacency index.*

*Permutation test*

A permutation test was conducted, with 1000 randomisations of the response. 58.6% of permuted models did not converge, so a Z-score was unable to be extracted. The two-sided permutation p-values supported the significance of all the coefficients which were found to be significant in the original model (Figure A8). However, the coefficient New:Day which was found to be non-significant in the original model was significant based on the permuted p-values. As this effect being significant would only strengthen the conclusions of our research, we chose to err on the side of caution, and continued to interpret this coefficient as non-significant based on the more conservative results of the original model.

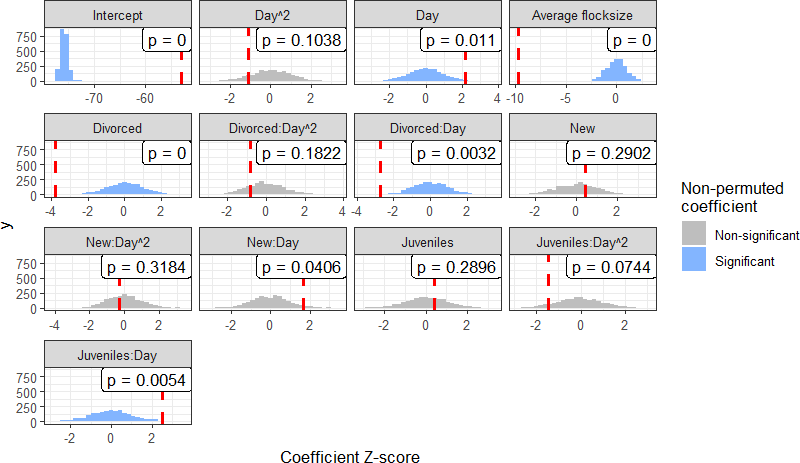

*Figure A8: The result of a permutation test of a model of the effect of pair status and day on visit adjacency index. Non-converged models with NA Z-score values are not included.*

*Coefficient table*

*Table A9: Fixed effect coefficient estimates from a binomial GLMM of the effect of pair status and day on visit adjacency index. Significant coefficients are indicated in italics.*

| **Coefficient** | **Estimate** | **Standard**  **error** | **p-value** | **Permuted**  **p-value** |
| --- | --- | --- | --- | --- |
| *Intercept* | *-2.93* | *0.06* | *<0.001* | *0* |
| Pair status New | 0.03 | 0.07 | 0.68 | 0.29 |
| Pair status  Juveniles | 0.03 | 0.08 | 0.67 | 0.29 |
| *Pair status*  *Divorced* | *-0.49* | *0.13* | *<0.001* | *0* |
| *Day* | *0.05* | *0.02* | *0.03* | *0.01* |
| Day^2^ | -0.03 | 0.03 | 0.27 | 0.10 |
| *Average*  *flocksize* | *-0.18* | *0.02* | *<0.001* | *0* |
| Pair status  New*Day | 0.06 | 0.04 | 0.102 | 0.04 |
| *Pair status*  *Juveniles*Day* | *0.10* | *0.04* | *0.013* | *0.01* |
| *Pair status*  *Divorced*Day* | *-0.23* | *0.08* | *0.006* | *0.003* |
| Pair status  New*Day^2^ | -0.01 | 0.03 | 0.757 | 0.32 |
| Pair status  Juveniles*Day^2^ | -0.06 | 0.04 | 0.150 | 0.07 |
| Pair status  Divorced*Day^2^ | -0.07 | 0.08 | 0.412 | 0.18 |

While the model with quadratic terms outperformed the linear model in model selection, none of the quadratic terms are significant in the final model. The non-quadratic model coefficient table is provided here for comparison.

*Table A10: Fixed effect coefficient estimates from a binomial GLMM of the effect of pair status and day on visit adjacency index. Significant coefficients are indicated in italics.*

| **Coefficient** | **Estimate** | **Standard error** | **p-value** |
| --- | --- | --- | --- |
| *Intercept* | *-2.96* | *0.05* | *<0.001* |
| Pair status New | 0.02 | 0.06 | 0.74 |
| Pair status Juveniles | -0.03 | 0.06 | 0.70 |
| *Pair status Divorced* | *-0.55* | *0.11* | *<0.001* |
| *Day* | *0.06* | *0.02* | *0.008* |
| *Average flocksize* | *-0.18* | *0.02* | *<0.001* |
| Pair status  New*Day | 0.06 | 0.04 | 0.120 |
| *Pair status*  *Juveniles*Day* | *0.08* | *0.04* | *0.029* |
| *Pair status*  *Divorced*Day* | *-0.20* | *0.08* | *0.008* |

**A.2.6	Preferred partner by time**

*Model selection*

*Table A11: AICc model selection results for a model of partner preference by pair status and experimental day. The model used in analysis is italicised.*

| **Model**  **number** | **Fixed effects** | **Random**  **effects** | **Zero-**  **inflation** | **AICc** | ∆ **AICc** |
| --- | --- | --- | --- | --- | --- |
| ***1*** | *Pair status + Day*  *+ Day*^2^ *+ Pair status*Day + Pair status*Day*^2^ *+ Sex* | *Year +*  *Pair ID* | *-* | *7357.12* | *0* |
| **2** | Pair status + Day  + Pair status*Day  + Sex | Year +  Pair ID | - | 7388.70 | 31.58 |

*Model residuals*

The residuals for the partner preference model are shown below (Figure A9).The Kolmogorov-Smirnov test shows significant deviation - however, this test is very sensitive to deviation at high sample sizes, so a permutation test was not conducted.

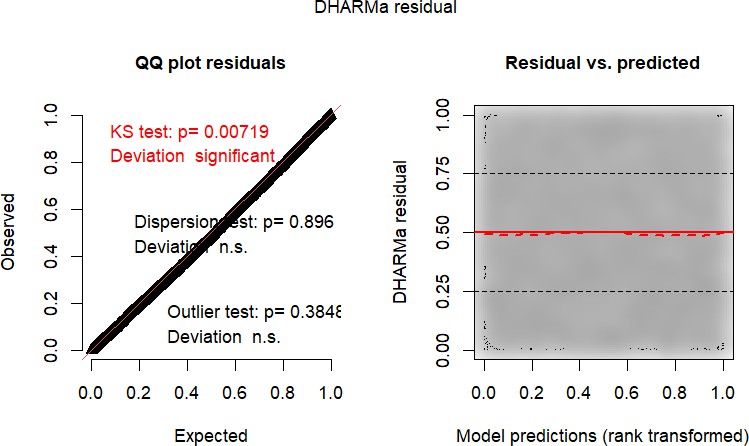

*Figure A9: The model residuals (estimated with DHaRMa) from a binomial GLMM of the effect of pair status and day on partner preference.*

*Coefficient table*

*Table A12: Fixed effect coefficient estimates from a binomial GLMM of the effect of pair status and day on partner preference. Significant coefficients are indicated in italics.*

| **Coefficient** | **Estimate** | **Standard error** | **p-value** |
| --- | --- | --- | --- |
| *Intercept* | *-1.38* | *0.13* | *<0.001* |
| *Pair status New* | *-0.76* | *0.17* | *<0.001* |
| *Pair status*  *Juveniles* | *-1.09* | *0.18* | *<0.001* |
| *Pair status Divorced* | *-2.84* | *0.41* | *<0.001* |
| *Day* | *0.05* | *0.38* | *<0.001* |
| *Day*^2^ | *-0.13* | *0.06* | *0.02* |
| Sex Male | -0.09 | 0.06 | 0.14 |
| *Pair status*  *New*Day* | *0.27* | *0.09* | *0.002* |
| *Pair status*  *Juveniles*Day* | *0.71* | *0.11* | *<0.001* |
| *Pair status*  *Divorced*Day* | *-0.67* | *0.23* | *0.003* |
| *Pair status*  *New*Day*^2^ | *-0.23* | *0.09* | *0.012* |
| Pair status  Juveniles*Day^2^ | -0.08 | 0.10 | 0.455 |
| Pair status  Divorced*Day^2^ | 0.21 | 0.25 | 0.389 |

**A.3	Non-pair associates**

As an additional analysis we also investigated how a bird’s winter association and visit adjacency scores with its partner compared to the same scores with its network associates. For each focal bird from the analysis, we found each bird that it had been observed sharing a flocking event with and then estimated winter association score and visit adjacency index using the same methods as in the other models. These estimates were then modelled, with methods and results presented below.

**A.3.1	Methods**

Winter association score was modelled using a generalised linear mixed model with a binomial response distribution. Pair type (original or associate) was used in interaction with pair status as a predictor, with year, pair, and a 1:n overdispersion term as random effects. A zero-inflation term was included, which was allowed to vary by pair type, as the selection of associates using shared flocking events meant that zeroes were only possible for ’original’ pairs. A permutation test was conducted to check for robustness of the results to violations of some residual assumptions.

Visit adjacency index was also modelled with a binomial GLMM, with pair status and type in interaction as fixed effects, as well as year, pair, and a 1:n overdispersion term as random effects. Average flock size was also included as a fixed effect predictor. AICc indicated no need to include a zero-inflation term. Visual analysis of the residuals using DHARMa showed no need for a permutation test.

**A.3.2	Results**

*Winter association score*

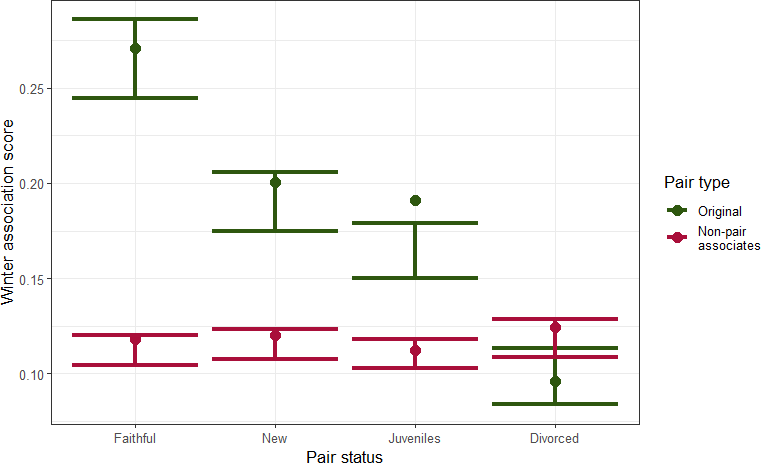

*Figure A10: The winter association score of birds with their breeding partner (’Original’) compared to their other network associates (’Non-pair associates’). Points represent means in the raw data, while error bars are the 95% prediction confidence interval calculated using ggeffects, averaged across random effect groups and conditioned on a zero-inflation term (Lüdecke, 2018).*

For faithful (coef=1.05*±*0.04, p=*<*0.001), new (Figure A10), and juvenile pairs (Figure A10), winter association scores with an individual’s breeding partner were significantly higher than winter association scores with their non-pair associates. For divorced pairs, there was no significant difference (Figure A10).

*Table A13: Fixed effect coefficient estimates from a binomial GLMM of the effect of pair type and pair status on winter association score. Significant coefficients are indicated in italics.*

| **Coefficient** | **Estimate** | **Standard error** | **p-value** |
| --- | --- | --- | --- |
| *Intercept* | *-1.02* | *0.05* | *<0.001* |
| *Pair status New* | *-0.43* | *0.06* | *<0.001* |
| *Pair status*  *Juveniles* | *-0.61* | *0.06* | *<0.001* |
| *Pair status Divorced* | *-1.21* | *0.09* | *<0.001* |
| *Non-pair associates* | *-1.05* | *0.04* | *<0.001* |
| *Pair status*  *New*Non-pair associates* | *0.46* | *0.05* | *<0.001* |
| *Pair status*  *Juveniles*Non-pair associates* | *0.59* | *0.05* | *<0.001* |
| *Pair status*  *Divorced*Non-pair associates* | *1.26* | *0.08* | *<0.001* |

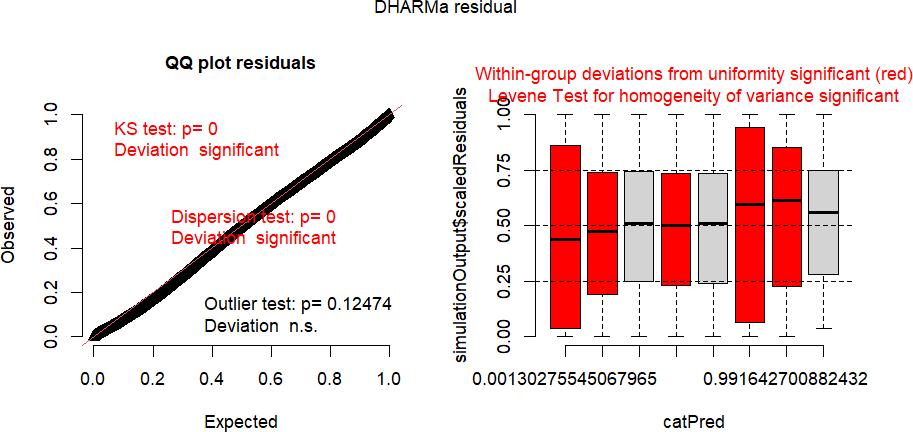

*Figure A11: The residuals for a binomial GLMM model of the effect of pair type and pair status on winter association score. Residuals calculated using the DHARMa package (Hartig, 2022).*

The model failed some checks of residual assumptions. While the likelihood of failing these checks increases at high sample sizes (n=20,418), a permutation test was conducted to be certain of model conclusions, following the methodology described in A.2.1. As shown in A12, the permutation test results concur with the results from the original model.

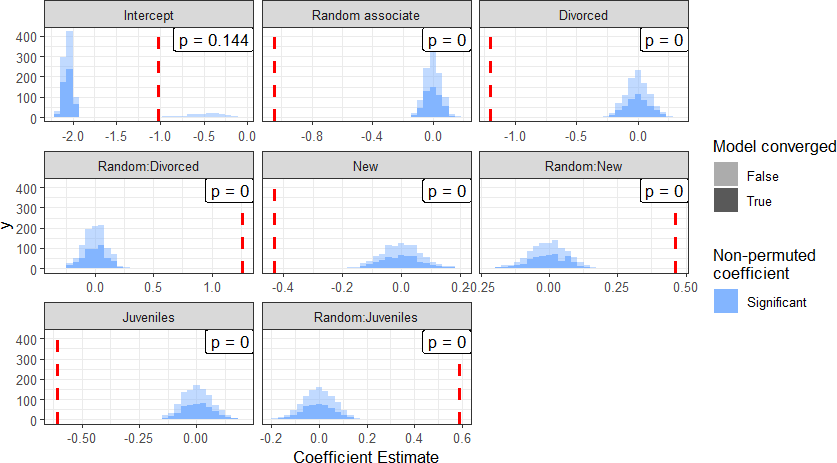

*Figure A12: The results of a permutation test for the coefficients in a binomial GLMM of the effect of pair type and pair status on winter association score.*

*Histograms represent the results from 1000 models of a randomly permuted dataset, while the red line represents the result from the original model.*

*Visit adjacency index*

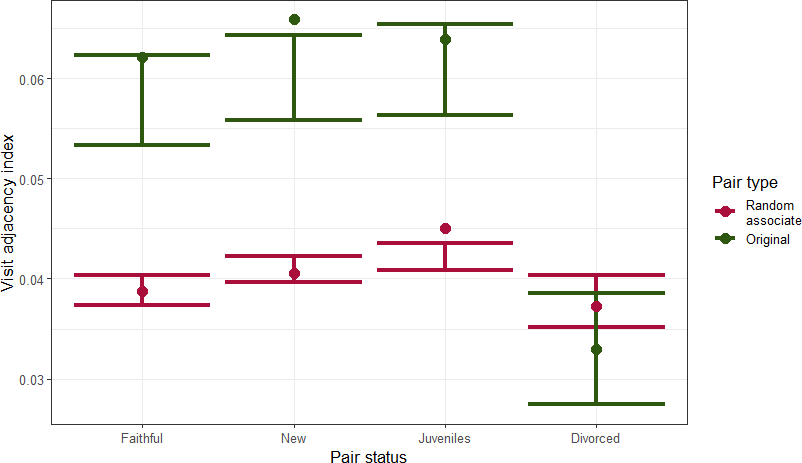

*Figure A13: The visit adjacency index of birds with their breeding partner (’Original’) compared to their other network associates (’Non-pair associates’). Points represent means in the raw data, while error bars are the 95% prediction confidence interval calculated using ggeffects, averaged across random effect groups and conditioned on a zero-inflation term (Lu¨decke, 2018).*

For faithful (coef=-0.42*±*0.04, p=*<*0.001), new (Figure A13), and juvenile pairs (Figure A13), visit adjacency index with an individual’s breeding partner was significantly higher than visit adjacency index with their non-pair associates. For divorcing pairs, there was no significant difference (Figure A13).

*Table A14: Fixed effect coefficient estimates from a binomial GLMM of the effect of pair type and pair status on winter association score. Significant coefficients are indicated in italics.*

| **Coefficient** | **Estimate** | **Standard error** | **p-value** |
| --- | --- | --- | --- |
| *Intercept* | *-2.70* | *0.06* | *<0.001* |
| Pair status New | 0.04 | 0.06 | 0.482 |
| Pair status Juveniles | 0.06 | 0.06 | 0.360 |
| *Pair status Divorced* | *-0.59* | *0.10* | *<0.001* |
| *Non-pair associates* | *-0.42* | *0.04* | *<0.001* |
| *Flocksizes* | *-0.0.1* | *0.001* | *<0.001* |
| Pair status  New*Non-pair associates | 0.01 | 0.05 | 0.793 |
| Pair status  Juveniles*Non-pair associates | 0.03 | 0.06 | 0.579 |
| *Pair status*  *Divorced*Non-pair associates* | *0.56* | *0.09* | *<0.001* |

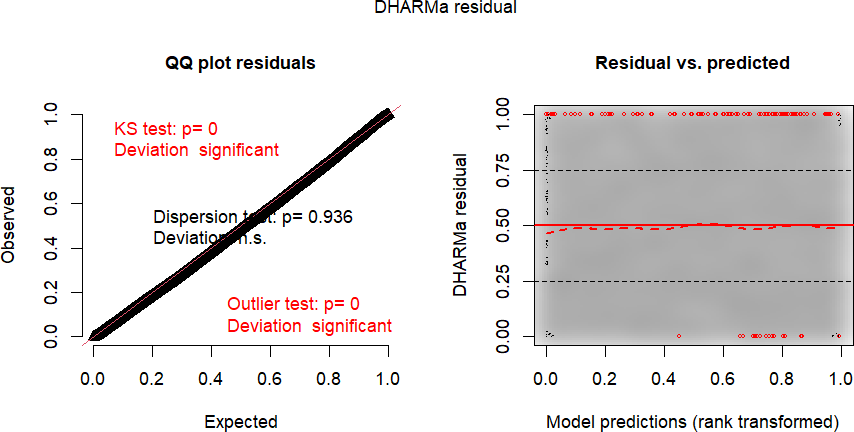

*Figure A14: The residuals for a binomial GLMM model of the effect of pair type and pair status on winter association score. Residuals calculated using the DHARMa package (Hartig, 2022).*

The model failed some residual assumption checks. However, a visual analysis suggests that this is due to the high sample size (n=20,388), so no permutation test was conducted.
